# Time of first contact determines cooperator success in a cross-feeding consortium

**DOI:** 10.1101/2024.05.13.593921

**Authors:** Rachel Los, Tobias Fecker, P.A.M. van Touw, Rinke J. van Tatenhove-Pel, Timon Idema

## Abstract

Microbial communities are characterised by complex interaction, including cooperation and cheating, which have significant ecological and applied implications. However, the factors determining the success of cooperators in the presence of cheaters remain poorly understood. Here, we investigate the dynamics of cooperative interactions in a cross-feeding consortium using individual-based simulations and an engineered *L. cremoris* toy consortium. Our simulations reveal first contact time between cooperators as a critical predictor for cooperator success. By manipulating the relative distances between cooperators and cheaters or the background growth rates, influenced by the cost of co-operation, we can modulate this first contact time and influence cooperator success. Our study underscores the importance of cooperators coming into contact with each other on time, which provides a simple and generalizable framework for understanding and designing cooperative interactions in microbial communities. These findings contribute to our understanding of cross-feeding dynamics and offer practical insights for synthetic and biotechnological applications.

## Introduction

Microorganisms often live together in surface-attached communities of many species, called biofilms. An early stage of the planktonic cell to biofilm lifecycle is a microcolonies, which make up the initial kernels that later grow into larger biofilms [1]. Studying the formation of these microcolonies provides valuable insight into biofilm development [2]. The species living in biofilms can form complicated networks of antagonistic, mutualistic, competing or cooperating interactions [3–5]. One of the ways species can cooperate is by cross-feeding, where both species produce metabolites or other essential compounds, that benefits the other [6]. The production of these compounds often comes at a cost for the producer in the form of extra expended energy, e.g. in the form of ATP. This creates opportunities for cheater species to exploit cooperators, by reaping the same benefits from the cross-fed compounds without contributing to the cooperation themselves [7]. Studying the interactions in bacterial systems can have implications for applications concerning biofilms in medical, biotechnological and industrial contexts [8–10]. Moreover, because mutualism and cooperative interactions can be observed in all branches of the phylogenetic tree, there is general interest in what makes this type of social behaviour evolutionary stable [7]. Due to their relative simplicity, microbial systems can be used as model systems for studying broader social behaviours in biology [11]. The interactions between microorganisms are in general complex, with many inter-dependent variables which are difficult to isolate [12]. An important factor on the dynamics of any collection of interacting species is spatial organisation [13–18]. This spatial structure plays a role in both the emergence and the maintenance of cooperator co-existence [19]. For example, spatially structured environments can promote clonal patching which is useful for intraspecies cooperation [20, 21] as well as pattern formation in multispecies colonies [16, 22].

In this paper we employ Individual-Based Modelling (IBM) to simulate a consortium of two cooperating species and a selfish cheater growing on a surface. By incorporating spatial structure and proximity in metabolic interactions in we attempt to uncover the factors that govern cooperative success in the face of cheating [23]. This will contribute to our understanding of population dynamics in cooperative biological systems.

## Results + Discussion

### Cooperator success is highly dependent on initial placement of particles

To explore the impact of local cross-feeding interactions on the development of a multi-species microcolony, we modelled interacting particles growing on a surface (Fig. 1a). We defined three different species, two cooperators A and B, and a cheater C (Fig. 1b). We made use of the fact that metabolic interactions have been shown to take place at very short distances and have the particles adjust their growth rate based on the number of their beneficial nearest neighbours [24, 25]. The strength of the cross-feeding interaction between the species is characterised by two main factors: the metabolic cost the cooperators pay to contribute to the cooperation, and the growth benefit they experience from this interaction. The cheaters don’t pay the cost but experience the same benefit (Fig. 1c). The particles, are spherocylindrical particles that grow and divide. We start each run with 10 of each species randomly distributed on a surface. We then let them grow till the colonies reach a size of 10^4^ particles.

**Figure 1.**
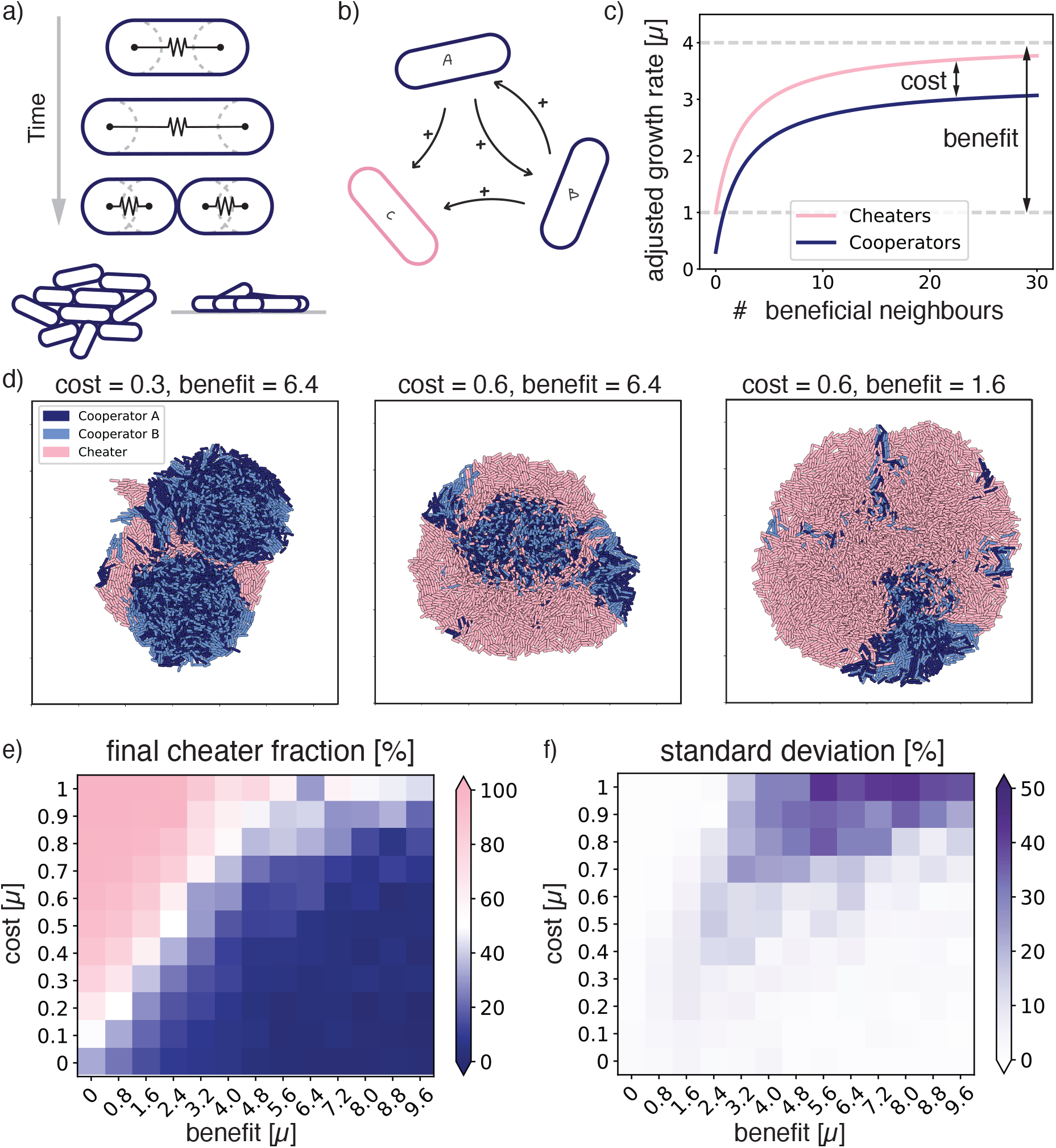
Effect of cost and benefit of cooperation on cooperator success in individual-based simulations. a) schematic representation of the individual-based-model. Spherocylindrical particles grow and divide, forming a 3D colony. b) The simulated consortium of two cooperators (A & B) which benefit each other and a cheater (C) which benefits from both A and B but does not reciprocate. c) Growth rate of both the cooperators and the cheaters, dependent on the number of beneficial neighbours (i.e., neighbours that provide a useful compound). The growth rate is affected by the cost and benefit of cooperation. d) Snapshots of the final 3D colonies for different combinations of cost and benefit. Area shown is always 130 *×* 130 µm^2^. e) Final cheater fractions for varying cost and benefit. Averages taken over 15 runs and in f) the standard deviation of those runs.

In order to understand how the strength of the local interactions shaped the development of the microcolony, we simulated growing colonies for varying costs and benefits and recorded the cheater fraction. Typical simulation snapshots of the final colonies are shown in Fig. 1d. The final cheater fractions averaged over 15 runs are shown in Figure 1e, the bottom left data point represents a situation with no cost and no benefit, which means that the particles are not interacting at all and their growth rate remains the same. As the cost of cooperation increases while the benefit stays zero, the final cheater fraction increases. In turn, increasing the benefit decreases the cheater fraction and favours the cooperators. Interestingly, there are not many intermediate values for the final cheater fraction, showing more switch-like behaviour than would be expected from a well-mixed system (Supplemental methods, Fig. S2).

Surprisingly, the simulations gave a large variation between runs with the same interaction parameters, particularly for higher costs and benefits (Fig. 1f). Here, we found standard deviations of up to 50% which indicates that the final cheater fraction can be anywhere between 0% and 100% for simulations with the same interaction input parameters. Since, the only difference between these runs was the initialisation of the 30 particles at the start of the run, the large variance had to be a result of the initial placement of the particles, which we explored next.

### First contact time determines cooperator success

In order to investigate how the initial placement of the particles leads to different final cheater fractions, we visually inspected the resulting colonies of the simulations. In general, we observed that colonies were fairly segregated, akin to clonal patches observed in 2D colonies [9]. The cooperator patches are often well mixed, with equal amounts of A and B, which we expect is due to a combination of nematic mixing and the mutualism between these species [20,26,27]. In different simulations with the same input parameters, we observed a varying number of these patches of mixed cooperators (Fig. S1). Because the cooperators depend on each other’s proximity to cross-feed, we speculated that the final cheater fractions might be determined by how likely it is for cooperators to meet.

In order to investigate this relation between the likelihood of meeting and final cheater fraction, we took all the runs and we recorded the first time point where an A particle comes into contact with a B particle. Plotting the final cheater fraction as a function of these first contact times for each value of the benefit, yielded clearly defined curve with comparatively little variance (Fig. 2a). Therefore, for a given benefit, the time of first contact was a more informative predictor for cooperator success than the cost of cooperation.

**Figure 2.**
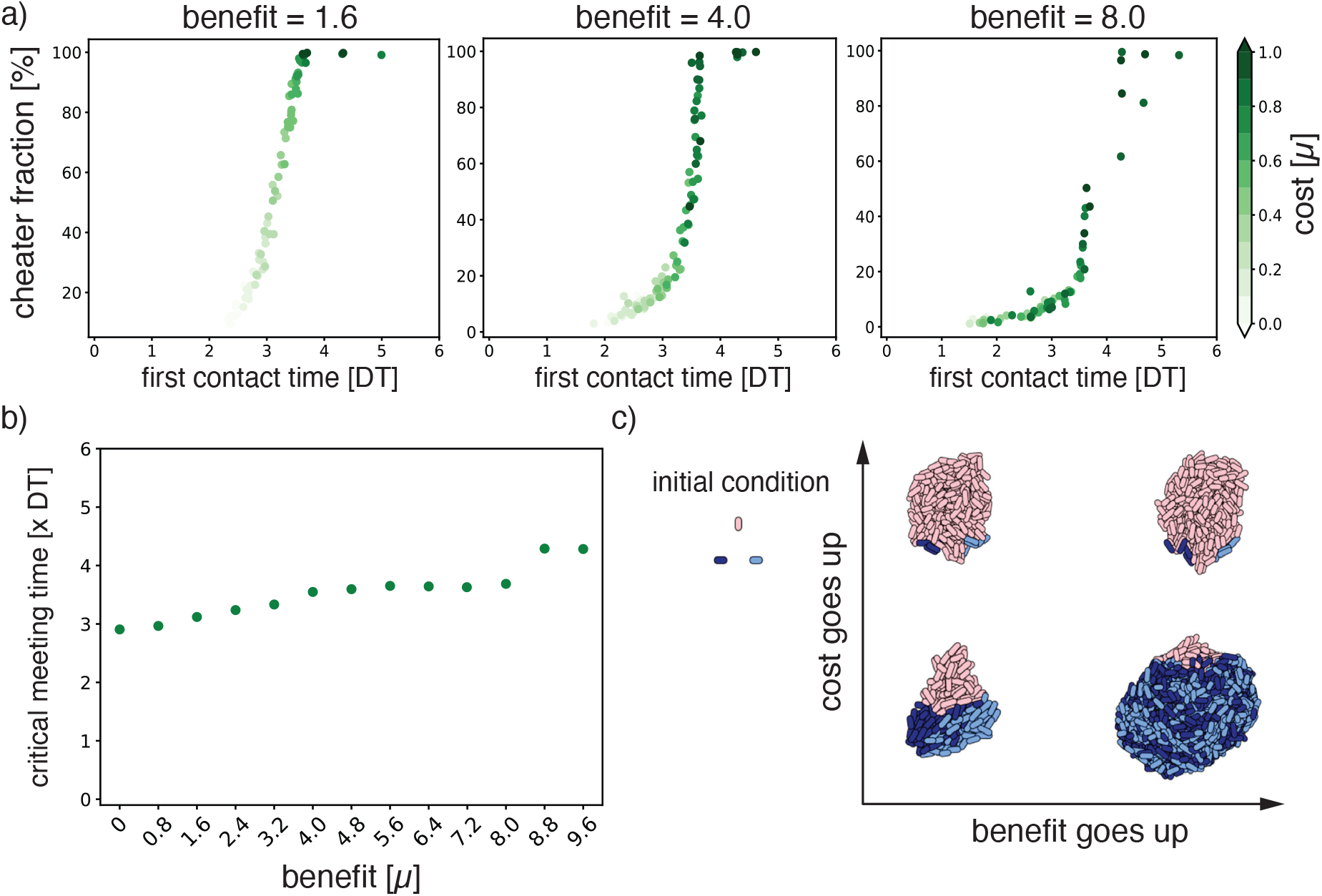
First contact time determines cooperator success. a) Final cheater fraction of single runs plotted as a function of first encounter time (measured in DT = 10^5^ simulation time steps) for three different benefits. Cost is denoted in green. b) Critical meeting time as a function of benefit. c) Toy system for a well defined initial condition of three particles positioned in a triangle and the resulting colonies, for the combinations of two values for the cost (0.2 and 0.7) and two values for the benefit (5.6 and 8.8).

This dependence on cooperators encountering each other is in line with earlier findings on the importance of co-localisation probabilities in a similar consortium growing in microdroplets [28]. Other work on two-species cooperator-defector consortia growing plates showed the importance of founder cell configuration [29]. Additionally, expansion-collision dynamics and initial distance between microorganisms have been shown to be important factors for cooperator-defector co-existence [30]. Although all these systems are not identical, they do point to a general principle in cooperative dynamics. To explore this further, for each value of the benefit, we determined a critical meeting time before which A and B need to make contact in order to outcompete the cheaters and make up *>* 50% of the final population (Fig. 2b). If the first contact time happened after this point, cheaters dominated the final colony of 10^4^ particles. The critical meeting time depended on the cooperator benefit, where a higher benefit allowed for a later critical meeting time. In this case, once first contact had been made, cooperators were able to make up for lost time by growing faster once they were together. Conversely, if the cooperator benefit was lower, the cooperators needed to meet earlier to outcompete the cheaters. It is important to note that first contact time is determined not only by the cost and benefit, but also the relative distance between the cooperators in the initial placement. To illustrate this, we visualised the outcome of simulations with a well-defined initial placement of the three particles (Fig. 2c). Here, the initial distance between the particles is always the same, and out of range from each other. The cost then affects the time it takes for the particles to traverse that distance since the cost affects the growth rate of the particles when they are on their own, i.e. their background growth. A higher cost causes A and B to have a lower background growth rate, so they are slower to reach each other. If they are too slow, as shown in the top-left, cheater particles grow in between them before they get the chance to make contact. In this case, having a higher benefit (top-right) will not help the cooperators as they never get to reap the rewards of their cooperation. If the cost is lower and A and B find each other before the critical meeting time (bottom-left), they can compete effectively with the cheaters. When the benefit becomes larger (bottom-right), they can significantly out-compete the cheaters once they meet. In our system of random initial distances between particles, this interplay between distance and background growth is what leads to the high variation in competitive outcome. From these simulation results, we propose that it is not just the initial distances between cooperators nor just the details of the cooperative interaction, but it is the first contact time that determines cooperator success in surface attached colonies.

### Experimental consortium of engineered *L. cremoris*

To test if the first contact time between cooperator species is also critical in a biological system, we attempted to translate our simulated consortium to a laboratory setting. We constructed a consortium of three *L. cremoris* strains (Fig. 3a, Table 1) growing on agar plates containing lactose as a carbon source and casein as an amino acid source. Cooperator A, NZ9000 Lac^+^ Glc^*−*^, can metabolize the galactose moiety of lactose, but exports the glucose moiety from the cell. The glucose can then be used by cooperator B, MG610, which cannot metabolise lactose. In return, MG610 expresses an extracellular enzyme to break down the casein into amino acids, which benefits A, which cannot metabolize the casein. Lastly, we engineered a cheater C, MG1363 GFP Lac^+^, by transforming a plasmid containing the genes necessary to metabolise lactose into MG1363 GFP. This constructed cheater can metabolise lactose and both resulting moieties, but, similar to A, lacks the enzyme to degrade casein. C also expresses GFP, which we use to determine the cheater fraction. When we grew A, B and C individually on agar plates, there was very little growth, which is what we expected from the designed cross-feeding interaction (Fig. 3b). When growing all the pairwise combinations, we observed that A+B massively outgrew the other combinations. From this we concluded that, as expected, these two species experience a strong mutual benefit from growing together. Similarly, we concluded that on plates with C+A and B+C, there is no such mutualism, as the total cell abundance, measured in amount of doublings, stalled. Furthermore, when we measured cheater fractions for these combined plates, we saw that C outgrows both A and B in a pairwise combination, pointing to a general advantage in growth rate that C has over both A and B (Fig. 3c). All in all, we concluded that our experimental consortium accurately represents a cross-feeding pair together with a cheater.

**Table 1:**
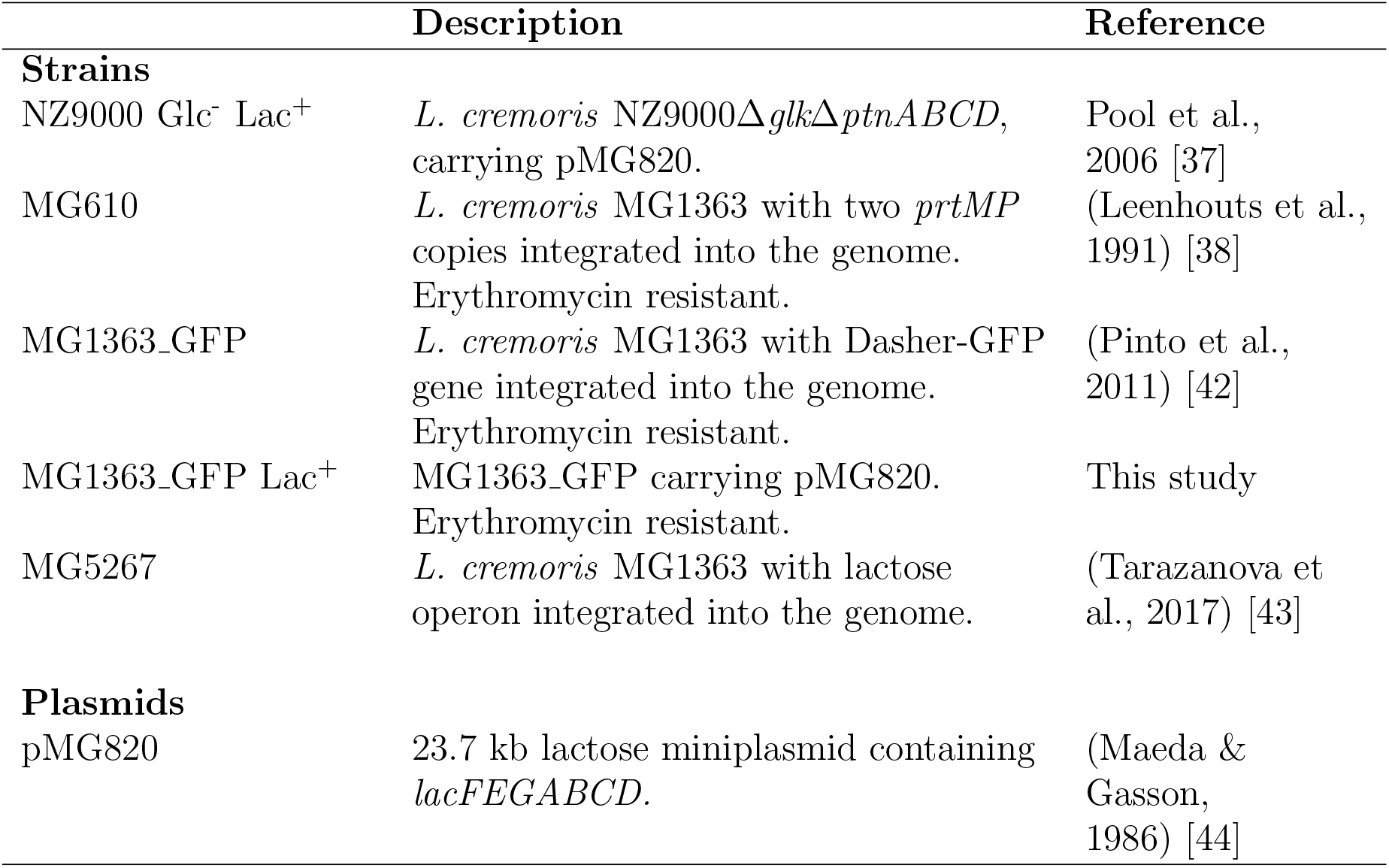
Strains and plasmids used in this study.

**Figure 3.**
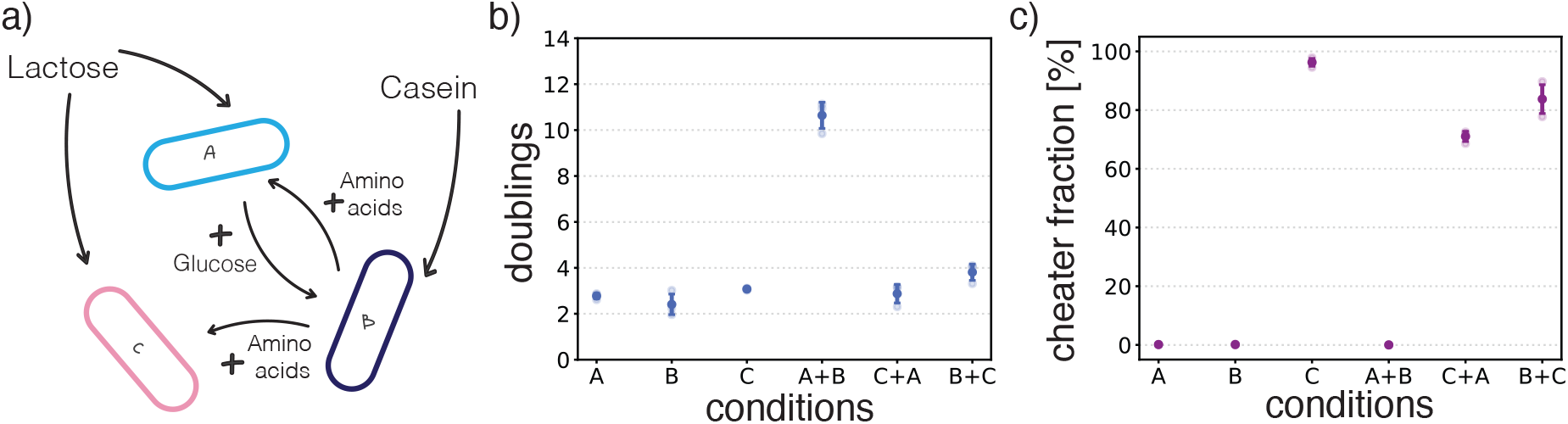
Engineered consortium on agar plates. a) *L. cremoris* consortium, consisting of cooperators A and B and cheater C. b) Control experiments showing the amount of doublings for single strain plates and pairwise combinations after three days incubation. c) Control experiments showing cheater fractions for all single strain plates and pairwise combinations. All conditions are measured in triplicate.

### Relative average distance between cooperators determines cooperator success in plate experiments

To investigate how the first contact time would affect our experimental system, we needed a way to modulate the distance between cooperators on the plates. Unfortunately, it is not so simple to closely control the initial placement or the respective growth rates of the microorganisms on plates. Therefore, we needed a different way to adjust the chance of cooperators meeting each other. Unlike our simulated cheater, the cheater in our experimental consortium is not symmetric in the sense that it does not benefit as much from A as it does from B. Since both A and C need B for their amino acid production, we argue that the relative difference between the distance between B and A (*r*_AB_) and the distance between B and C (*r*_CB_) sets the chance of cooperators meeting. In essence it’s a race between A and C, to reach B first.

To modulate the relative distances between A and B, and C and B, we adjusted the ratio of [C]/[A]. When [C]/[A] is small (*<* 1), there is more A on the plate than C. In this case, the chance of B finding itself close to an A is larger than B finding itself close to a C. If [C]/[A] is large (*>* 1), the reverse is true, so on average it is more likely for a B to be surrounded by cheaters (Fig. 4a). The relationship between [C]/[A] and the expectation value of the relative distance can be derived analytically (see SI, Fig. 4b).

**Figure 4.**
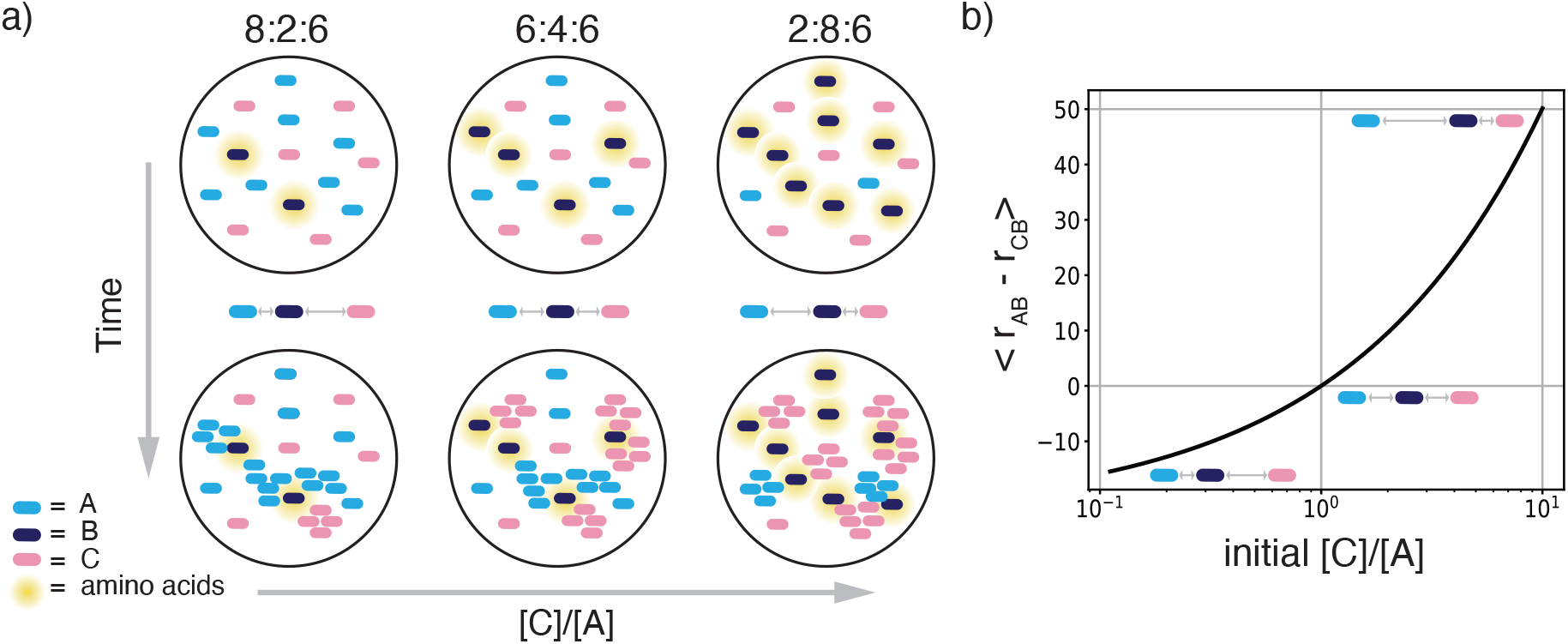
[C]/[A] ratio as a proxy for relative cooperator-cheater distances. a) Schematic of how changing the A:B:C ratios is expected to affect the first contact time between A and B. b) Relation between the expected average difference between the distance from B to A (*r*_AB_) and from B to C (*r*_CB_) to the [C]/[A] ratio. For the derivation, see Supplemental methods.

From our simulation results, we expected the final cheater fraction to rise with a higher relative distance between cooperators, as it would take longer for them to find each other. To test if this would also occur in our experimental consortium, we inoculated cells on plates in different A:B:C ratios, and after 3 days of incubation we washed the plates and measured the total cell count and the final cheater fraction (Fig. 5a & S3).Consistent with our expectations, a small [C]/[A] resulted in final cheater fractions of as low as 20%,where when the ratio was high, the final cheater fraction increased to almost 70%. Note that the total amount of cells after incubation was similar for all starting ratios (Fig. S3).

**Figure 5.**
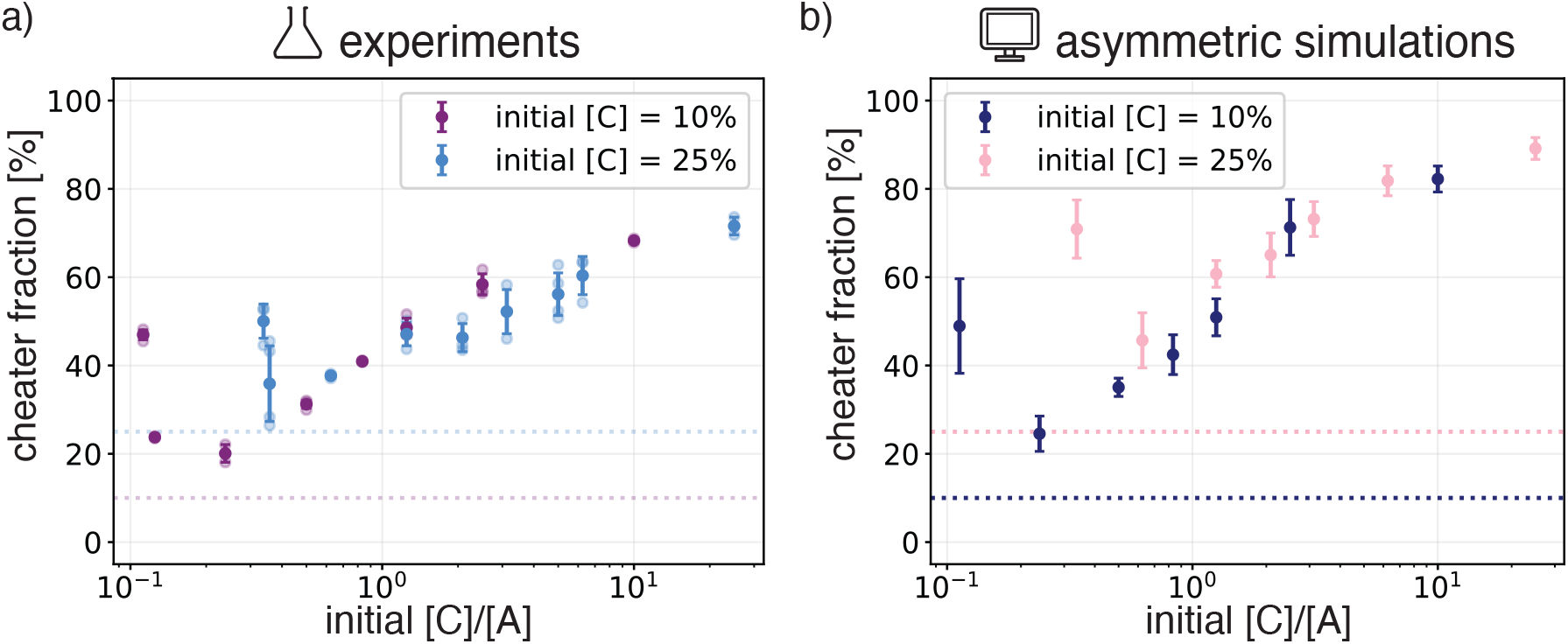
Final cheater fractions depend on initial A:B:C ratios in experiments and simulations. a) Final cheater fraction of plate experiments with different initial starting ratios of A:B:C, with a constant starting fraction of 10% C in purple and 25% C in blue. Error bars are standard deviation for 3 replicates. Doublings for these experiments can be found in Fig. S3. b) Final cheater fraction for simulations with the same starting ratios of A:B:C. Simulations are of an asymmetric interaction where A and B interact as before, but C only benefits from B. The cost and benefit for these runs are 0.5 and 2.4.

We show 2 sets of experiments, one where the initial cheater fraction is always 10% and one where it is 25%. Apart from the leftmost points of the respective curves (discussed below), they fall onto the same curve. From this we concluded that, regardless of initial cheater fraction, the relative abundance of A and C and therefore the relative average distance between cooperators determines cooperator success or failure in our experimental consortium.

### Too few nucleation sites result in higher cheater fractions

As shown in Fig. 5a, most data points fall onto the same curve. However, for the smallest [C]/[A] for both the 10% and the 25% curve, the final cheater fractions were higher than expected. These points correspond to A:B:C ratios of 89:1:10 and 74:1:25, respectively, so in both cases there is a minimal amount of B present. Because there is so much more A than C present, we had expected that B coming into contact with A would be inevitable, and therefore the final cheater fraction would be low. Instead, we measured the final cheater fraction at low B to be around 50%.

In order to generate a hypothesis of what happens for low amounts of B, we went back to our simulations. To better reflect the asymmetric interaction the cheater has with cooperators A and B in our experimental consortium, we adapted the model so that C would only benefit from B, while keeping the interaction between A and B the same. When we performed these asymmetric simulations for all the ratios we tested in the experiments, the simulations results were similar to the experimental data (Fig. 5a & b). Additionally, we indeed got the same sudden increase in cheater fraction for very low B. Following the trajectories of the composition of these colonies over time, we see that indeed, B always finds A (Fig. S4). However, because there is only one nucleation site for a cooperator patch, the development of the colony takes longer, and by the time A and B have grown to a significant size, the cheaters have already taken up a large part of the colony. This suggests that, next to meeting on time, there is also a minimal amount of nucleation sites necessary for cooperator success.

### Lowering the background growth of cooperators results in higher cheater fractions

The symmetric simulations demonstrated that not only the initial distance between cooperators determined cooperator success, but also how fast this distance could be traversed (Fig. 2c). This speed is set by the background growth of the particles. In the simulation, we can set the background growth of the cooperator particles by changing the cost of cooperation (Fig. 2c). A higher cost for cooperators means a lower background growth, which we expected to result in a shift of the curve to the left, where more of A, i.e. a smaller [C]/[A] ratio, would be needed to achieve the same final cheater fraction. Indeed, increasing the cost in the asymmetric simulations slightly, results in a shift to the left, shown as the blue data points in Fig. 6b.

**Figure 6.**
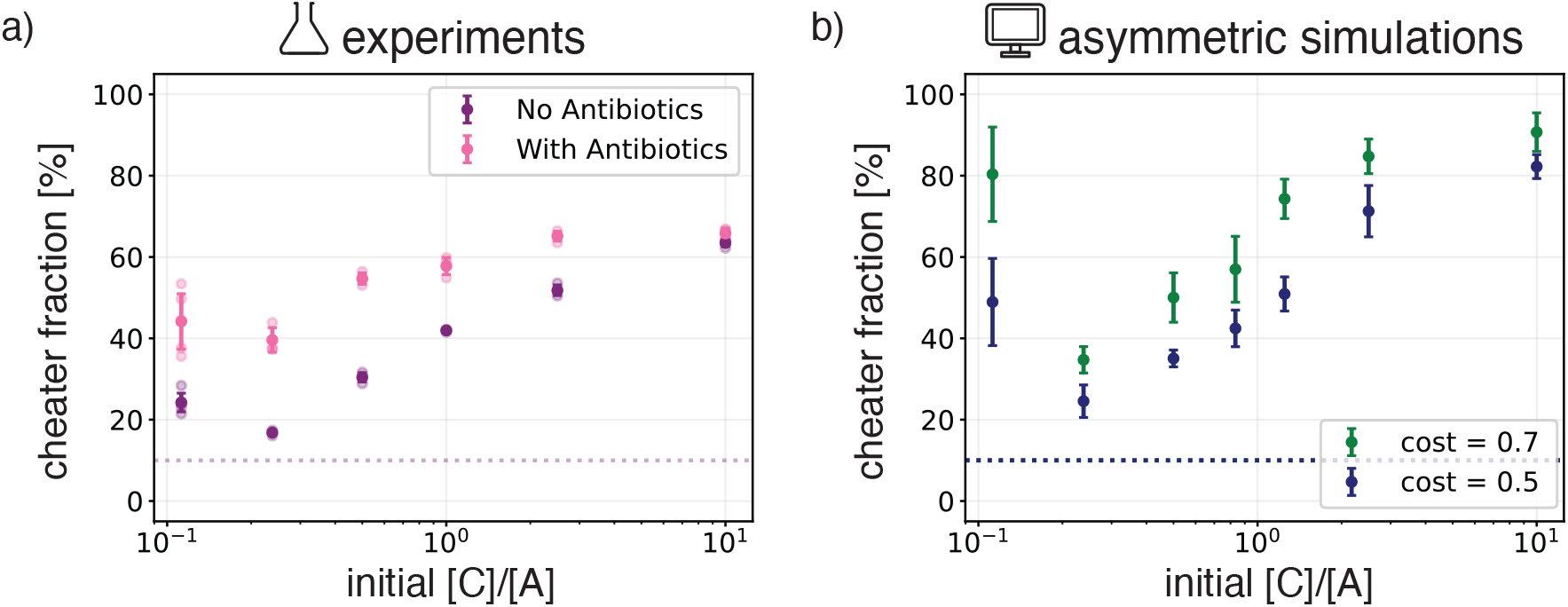
Higher cost of cooperation leads to higher cheater fractions. a) Final cheater fraction for experiments with different [C]/[A] initial ratios on plates containing 0.04 µg mL^*−*1^ erythromycin (in pink) and plates without erythromycin (in purple). b) Final cheater fraction for asymmetric simulations of different [C]/[A] initial ratios for two values of the cost. The benefit is 2.4 and all data points are averages and standard deviations of 5 runs.

Again, we attempted to test these results from the simulations experimentally. While the cost and benefit of cooperation could not be directly altered, we hypothesised that by selectively inhibiting A we could achieve a similar effect. To that end, we used erythromycin which is an antibiotic that only affects the growth of A as both B and C are resistant. We tested for a concentration of erythromycin that would only slightly inhibit the growth of A, which we found to be 0.04 µg mL^*−*1^ (Fig. S5). We then performed the same experiment for the 10% initial cheater fraction on both normal (untreated) plates and on plates containing the antibiotic (Fig. 6a). For the final cheater fraction on the antibiotic plates, we observed a shift to the left compared to the normal plates, where the same final cheater fraction on standard plates could be achieved on antibiotic plates with a lower ratio of [C]/[A].

Note that in the simulations, we increased the cost for both A and B, where in the experiments, the antibiotic only affects A. Regardless, we achieved good agreement between the experimental and simulated data. This demonstrates that increasing the cost of cooperation or decreasing the background growth of the cooperators results in higher cheater fractions. Together with our previous results, we propose that this is because the background growth sets the first contact time for the cooperators, which is the main determinant for cooperator success.

The general concept of first contact times being instrumental in cooperator success, suggests several strategies for cooperators to increase their survival rate. For instance, if cooperators could increase their affinity for each other in solution, forming mixed aggregates. These would then function as the kernels that establish colonies elsewhere, which would greatly improve their chance of survival [1]. Another option would be for cooperators to economise on the production of the cross-fed compound when growing on their own and using quorum sensing to only start production when encountering other cells [31, 32]. Both these strategies could also be explored in synthetic systems making use of specific adhesins and quorum sensing pathways [33, 34].

## Summary

Microbial collaboration is an abundant phenomenon with ecological and applied relevance, yet the factors contributing to cooperator success in the presence of cheaters are poorly understood. We set out to investigate the factors contributing to cooperator success in the presence of an exploiting cheater growing together on a surface. Individual-based simulations of a cross-feeding consortium consisting of three species indicated a strong influence of the initial placement of the microorganisms on the final outcome. Focusing in on the mechanisms, we demonstrated that first contact time was a better predictor for cooperator success than the value of the interaction parameters alone. We then showed how a combination of cost and initial placement together affect this first contact time.

We translated our simulations to an engineered *L. cremoris* toy consortium, consisting of two mutualistic strains and a cheater strain growing on agar plates. We show that by changing the relative distance between cooperators and cheaters by altering the starting ratios of A, B and C, we could directly influence cooperator success. We recreated our experimental findings by a simple adjustment to the model, making the cheater an asymmetric cheater, further showing how the average distance between cooperators is responsible for cooperator success.

Finally, we showed that, next to the relative distance between cooperators, the time it takes to traverse that distance affects the final cheater fraction as well. This time is set by the background growth, which is dependent on the cost of cooperation in the simulations. In the experiments we used antibiotics to selectively inhibit the background growth of A, giving good agreement with the simulations.

We conclude that in a cross-feeding cooperative interaction between strain A and strain B, the ability to find each other on time is the determining factor in cooperator success in the presence of a cheater strain.

Our findings provides better understanding of cross-feeding dynamics in surface attached communities, as well as an intuitive framework for designing and altering cross-feeding consortia in synthetic and biotechnological applications.

## Materials and Methods

### Individual based model

In our simulations we initialise spherocylindrical particles on a surface and let them grow, divide and interact with each other [35]. The mechanical interactions of the particles are the same for all species: in addition to steric repulsion between the particles, they have an attractive potential which makes them stick together and to the surface (see SI). The cross-feeding interaction between the different species is implemented by adjusting their growth rates based on their immediate environment (Fig. 1c) [24, 25]. In particular, for every growth step, we count for each particle how many beneficial neighbours they have in their immediate vicinity and use this number as input for a scaled Monod equation:

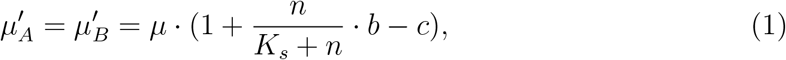

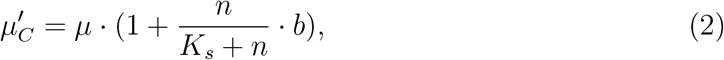

which determines the growth rate of the particle, *µ*′. Here, *n* is the number of beneficial neighbours, *K*_*s*_ is the Hill coefficient, which we set to 2.5 as this is the average number of beneficial direct neighbours in a well-mixed colony of 10^4^ particles, *µ* is the background growth of the cheaters, i.e., the growth rate of an isolated cheater particle. Both benefit, *b*, and cost, *c*, are given as fractions of *µ*, where *b* sets the maximal growth rate a particle can achieve and *c* gives a downward shift which can have any value between 0 and 1.

### Cross-feeding simulations

We implement a symmetric cross-feeding interaction by having species A and B increase their growth rate depending how many of their neighbours are B and A, respectively. C increases its growth rate based on the minimum number of A and B in its vicinity. At the beginning of the simulation, 10 particles of each species are randomly initialised on a 50 *×* 50 µm^2^ patch of surface. We grow the system to a final size of 10^4^ particles.

### Asymmetric simulations

For the asymmetric case that resembles our experimental consortium, the interactions remain the same except that for the growth rate of C, *n* is taken to be the number of B particles in the neighbourhood. Additionally, 100 particles are initialised in different A:B:C ratios, and the system is grown to a final size of 7 *·* 10^3^ particles.

### Microbial strains and growth conditions

Strains and plasmids used in this study are listed in table 1. All strains were grown on CDMpc, described by Price *et al*. [36] at 30°C in a stationary incubator. We used glucose and lactose as carbon sources, in the concentrations as indicated. Where indicated, the strains were grown on CDMpc_cas_, containing 0.2 wt% casein sodium salt (from bovine milk, #C8654, Sigma Aldrich, Saint Louis, MO, USA) instead of amino acids. To prepare agar plates, liquid medium was supplemented with 2 wt% BD Difco™ Bacto™ Agar (BD, NJ, USA).

*L. cremoris* NZ9000 Glc^-^ Lac^+^ [37], *L. cremoris* MG5267 and MG1363 GFP Lac^+^ (this study) were precultured in 25 mL CDMpc + 0.09 wt% lactose, *L. cremoris* MG610 [38] and MG1363 GFP [39] were precultured in 25 mL CDMpc + 0.09 wt% glucose. Freezer stocks were prepared by growing the strains in CDMpc with the appropriate carbon source and storing the stationary culture at -80°C with 20 vol% glycerol. Where indicated, erythromycin (Sigma-Aldrich, 856193, Saint Louis, MO, USA) was added at the indicated concentration. For the isolation of pMG820 and transformation into MG1363 GFP, M17 medium was used (from powder, Thermo Fisher Scientific, MA, USA).

### Isolation of pMG820 and transformation into MG1363 GFP

To construct MG1363 GFP Lac^+^, pMG820 was isolated from 5 mL of a stationary NZ9000 Glc^-^ Lac^+^ culture, cultivated in M17 + 0.5 wt% lactose, as follows. The culture was centrifuged using a Sorvall ST40 centrifuge (Thermo Fisher Scientific, MA, USA) (10 min, 5000g), and resuspended in 30 mM Tris-Hcl pH 8, 3 mM MgCl_2_, 25 wt% sucrose and 2 mg/mL lysozyme (from egg white, 10837059001 Roche, Basel, Switzerland) and incubated for 30 min at 37 °C. Afterwards, the plasmid was isolated using the GeneJET Plasmid Miniprep Kit (Thermo Fisher Scientific, MA, USA). Once obtained, the plasmid was transformed into MG1363 GFP as described by Wells et al. [40] with the following adaptations. The MG1363 GFP cells were precultured in 50 mL M17 broth with 17 wt% (0.5 M) sucrose, 2.5 wt% glycine and 0.5 wt% glucose at 30 °C. After overnight incubation, they were centrifuged (6000 g, 20 min) and washed with 400 mL 17 wt% (0.5 M) sucrose, 10 wt% glycerol (4°C), spun down and resuspended in 200 mL 17 wt% (0.5 M) sucrose, 10 wt% glycerol + 50 mM EDTA (4°C). After incubating on ice for 15 min and spinning down (6000g, 10 min) the cells were washed again as described above and resuspended in 4 mL 17 wt% (0.5 M) sucrose, 10 wt % glycerol (4°C). 40 µL of the cell solution with 1 µL DNA (100 ng/µL in Tris-Buffer) was added to a chilled cuvette and pulsed using a Bio-rad Genepulser (Bio-Rad, CA, USA) for 5.7 ms (2000 V, 25 µF, 200 Ω), after which 1 mL M17 broth with 17 wt% (0.5 M) sucrose, 2.5 wt% glycine, 0.5 wt% glucose, 20 mM MgCl_2_ and 2 mM CaCl_2_ was added. The solution was added to M17 agar plates containing 0.5 wt% lactose. After incubation for 48 h, colonies were picked and restreaked.

### Growth rate determination

To check the phenotype of *L. cremoris* MG1363 GFP Lac^+^, we tested whether the strain was fluorescent, and whether the growth rate was similar to the growth rate on lactose of *L. cremoris* MG5267, a lactose-positive *L. cremoris* strain, and similar on glucose to its ancestor strain MG1363. Fluorescence of the strain was confirmed using flow cytometry (Accuri C6, BD, NJ, USA). To prepare the cells for inoculation for the growth rate determination, MG1363 GFP Lac^+^ was precultured in CDMpc on both glucose and lactose, and MG5267 and MG1363 were precultured as described above. After 16 h, the OD_660_ was determined using a Jenway 7200 Spectrophotometer (Cole-Palmer, Stone, United Kingdom) and the cells were inoculated in duplicates in 30 mL of the same medium as in the preculture at an OD_660_ of 0.01. To determine the growth rate, the OD_660_ was measured every hour (Fig. S7).

### Co-cultivation on plates

*L. cremoris* MG1363 GFP Lac^+^, NZ9000 Glc^-^ Lac^+^ and MG610 were precultured as described above, spun down (8000g, 15 min), washed in sterile PBS and resuspended in PBS to a final concentration of 3x10^7^ cells / mL, as determined by flow cytometry (Accuri C6, BD, NJ, USA). To make the cell mixtures, cells were added together to the indicated cell concentration and strain ratio. Of the cell mixtures, 100 µL was added to plates containing CDMpc_cas_.The plates were incubated for 90 h at 30 °C. Afterwards, the cells were removed from the plate by adding 2 mL sterile PBS and spreading it using a sterile spreader, as described in [41]. The cell suspension was removed from the plate, and the cell concentration and cheater abundance was determined using flow cytometry (Accuri C6, BD, NJ, USA). To selectively lower the growth rate of cooperator A, erythromycin was added to the agar plates at the indicated concentrations.

## Author contributions statement

R.L. designed the research. R.L and T.I. designed the simulations. R.L. and P.v.T. performed simulations and analysed the data. R.L. and T.F. performed experiments and analysed the data. T.F. and R.v.T. designed the experiments. R.v.T. and T.I. supervised the research. R.L. wrote the manuscript with input from all authors.

## Data availability

All experimental data are provided in the article and the online supplementary material. The simulation data can be found on the 4TU repository, at https://doi.org/10.4121/1d711dde-3128-4b14-9a18-444e195361d6.v1.

## Acknowledgments

We thank Sagarika Bangalore Govindaraju for her support with the experiments. We thank Felix Frey for reviewing the manuscript.

