## Supplemental Methods and Figures for "Time of first contact determines cooperator success in a cross-feeding consortium"

### Supplemental Information for "Time of first contact determines cooperator success in a cross-feeding consortium"

May 13, 2024

1 Department of Bionanoscience, Kavli Institue of Nanoscience, TU Delft, The Netherlands.
2 Department of Biotechnology, TU Delft, The Netherlands.

### 1 Supplemental methods

#### Individual-based simulation details

Each particle is modelled as a spherocylinder that consists of two spheres of diameter, $D$, connected by a spring with a rest length $L$ (Fig. 1a). This rest length is incrementally increased with growth rate $\mu$ until it reaches the division length $L_d$. The particle then splits up into two daughter particles. Here, a tiny bit of noise is added in the orientation to make sure that the particles don't grow in a perfect line. For each particle pair that is within interaction range, $R_{int}$, of each other, the repulsive or attractive force is determined. First, the shortest distance between the particles is determined. If it is negative, i.e. there is overlap between the particles or the surface, the repulsive force is calculated as a spring force with spring constant $k_r$, scaling with the amount of overlap $d$,

$$F(d) = -k_r \cdot d. \tag{1}$$

This force is then distributed along the backbone inversely proportional to where along the backbone the overlap occurred.

Particles also experience an attractive force between each other and with the surface. The magnitude of the particle adhesion force depends on the distance $r$ between the centres of the particles and their relative orientations in the following way:

$$F(r, \theta, \phi) = F_{PP} \cdot \left( 4 \left( \frac{0.95D}{r} \right)^5 - 4 \left( \frac{0.95D}{r} \right)^9 \right) \cdot \cos\theta \cdot \cos\phi, \qquad \text{for } r > D, \tag{2}$$

where $D$ is the diameter of the particles, $F_{PP}$ determines the amplitude of the attractive forces, and $\theta$ and $\phi$ are the angles between the backbones of the two particles. The resulting force is then distributed over the two end-points of the particle inversely related to where the closest point between the particles is situated along the length of the particle.

The attractive force to the surface is implemented similarly, but $r$ is now the distance from the center of the particle to the surface.

$$F(r, \theta) = F_{PS} \cdot \left( 4 \left( \frac{0.95R}{r} \right)^5 - 4 \left( \frac{0.95R}{r} \right)^9 \right) \cdot \cos\theta, \qquad \text{for } r > R, \tag{3}$$

where $R$ is the radius of the particles or half the diameter, and $\theta$ is the angle the particle makes with the surface. Again the force is distributed over the two spheres that make up the particle, inversely proportional to the distance of each end point to the surface. Consequently, the particles experience both a force and a torque due to the attractive and repulsive interactions.

Every time step all the forces that the particles undergo are calculated and they are moved accordingly. We grow all the particles once every 5 time steps. We grow the system to a set amount of particles and then count how many of those particles are cheaters. Values for all the parameters used can be found in Table 1.

| Parameter | Explanation | Value |
|---|---|---|
| $k_{int}$ | spring constant of the internal spring | $0.1\ \mathrm{N\,m^{-1}}$ |
| D | particle diameter | 1 µm |
| $L_d$ | length of the spring at division | 4 µm |
| $k_r$ | spring constant of the overlap potential | $0.2\ \mathrm{N\,m^{-1}}$ |
| $F_{PS}$ | particle-surface adhesion prefactor | $10^{-5}$ N |
| $F_{PP}$ | particle-particle adhesion prefactor | $10^{-5}$ N |
| DT | division time for a particle growing with growth rate $\mu$ | $10^5$ ts |
| $R_{int}$ | interaction range for which mechanical interactions are calculated | $2 \cdot L_d$ |
| $\mu$ | background growth rate for a cheater particle | $0.0002\cdot\ \mu\mathrm{m}(5 \cdot \mathrm{ts})^{-1}$ |
| $K_s$ | Hill coefficient for the growth rate adjustment | 2.5 |

Table 1: Explanation and values used for the simulation parameters.

#### Well-mixed model of costly cooperation

In order to show the effect of spatial structure on our simulated consortium, we compare it to a well-mixed system. For this we solve the following system of equations for the evolution of populations of interacting species A, B and C:

$$\frac{dA}{dt} = (1 - c + H(B, N, b))\, A, \tag{4}$$

$$\frac{dB}{dt} = (1 - c + H(A, N, b))\, B, \tag{5}$$

$$\frac{dC}{dt} = (1 + H(\min(A, B), N, b))\, C. \tag{6}$$

$$\tag{7}$$

Here $A$, $B$, and $C$ are the size of the population of the corresponding strain. $N$ is the total size of the population. The growth of the populations depends on $c$, which is the cost of cooperation, and $H$, which calculates the benefit depending on the ratio of the cooperative species as follows:

$$H(n, N, b) = \frac{\frac{n}{N} \cdot \bar{n}}{K_s + \frac{n}{N} \cdot \bar{n}} \cdot b, \tag{8}$$

$$\tag{9}$$

where $n$ is the amount of beneficial particles in the total population and $N$ is the size of the total population. Additionally, $b$ is the benefit of cooperation, $K_s$ is the Hill coefficient which is set to 2.5, and $\bar{n}$ is the average number of neighbours a particle has, which is set to 7.5.

The initial conditions are the same as for the simulations, so 10 of each strain, the cheater fraction is measured when a population size of $10^4$ particles is reached. The results are summarised in Fig. 2.

#### A:B:C ratios and how they relate to effective distance between cooperators

If we look from the perspective of a B particle on a plate, the amount of particles that we find within radius, $r$, away from that particle follows a Poisson distribution,

$$P(k) = \frac{\lambda^k \cdot e^{-\lambda}}{k!}, \text{ with } \lambda = \pi r^2 \sigma. \tag{10}$$

Here, $k$ is the amount of particles and $\lambda$ is the expectation value of the amount of particles in a circle of radius $r$, based on the density of the particles $\sigma$. Because we are interested in what the first encounter will be, we want to compare the radii for the first encounter with an A particle and the one with a C particle. The probability of having at least one A particle within a radius, $r$, is given by,

$$\begin{aligned} P(A) &= 1 - P(0), \\ &= 1 - e^{-\pi r^2 \sigma_A}. \end{aligned} \tag{11}$$

From this we can get a probability density function,

$$\begin{aligned} f(r) &= \frac{\mathrm{d}}{\mathrm{d}r} P(A), \\ &= 2\pi r \sigma_A \cdot e^{\pi r^2 \sigma_A}, \end{aligned} \tag{12}$$

which we can then use to find an expectation value of what the distance is that a B will encounter its first neighbour of type A.

$$\begin{aligned} \langle r_{AB} \rangle &= \int_0^\infty r \cdot 2\pi r \sigma_A \cdot e^{\pi r^2 \sigma_A} \,\mathrm{d}r \\ &= \frac{1}{2\sqrt{\sigma_A}}. \end{aligned} \tag{13}$$

Similarly, the first neighbour of type C is expected to be found at,

$$\langle r_{CB} \rangle = \frac{1}{2\sqrt{\sigma_C}}. \tag{14}$$

When we subtract the two we get the expected relative distance which denotes who is expected to be further away from B, A or C:

$$\langle r_{AB} - r_{CB} \rangle = \frac{1}{2\sqrt{\sigma_A}} - \frac{1}{2\sqrt{\sigma_C}}. \tag{15}$$

The particle density at inoculation, $\sigma$, can be easily calculated, as it is always the same, namely $3 \cdot 10^6$ cells on a petri dish with a diameter of 9 cm. This gives us an overall initial particle density of

$$\sigma = \frac{3 \cdot 10^6}{\pi (4.5 \cdot 10^4)^2}, \tag{16}$$

$$= 0.00465\ \mu\mathrm{m}^{-2} \tag{17}$$

The densities of A and C are then simply the fractions of those particles times this overall density. $\langle r_{AB} - r_{CB} \rangle$ is plotted for all [C]/[A] ratios used in the 10% cheater fraction experiments in Fig. 4b.

| **Strain** | **Carbon source** | **$R^2$ rep. 1** | **$R^2$ rep. 2** | **$\mu_{max}$ (1/h) ± SD** |
|---|---|---|---|---|
| MG1363-GFP ($Lac^+$) | 0.09 wt% glc | 0.997 | 0.998 | 0.67 ± 0.00 |
| MG1363-GFP | 0.09 wt% glc | 0.997 | 0.998 | 0.57 ± 0.01 |
| MG1363-GFP ($Lac^+$) | 0.09 wt% lac | 0.996 | 0.999 | 0.66 ± 0.00 |
| MG5267 | 0.09 wt% lac | 0.998 | 0.996 | 0.64 ± 0.00 |

Table 2: Goodness-of-fit of all extracted growth curves as determined by the $R^2$ of the $\ln(OD_{660})$ over time and resulting $\mu_{max}$ ± SD (n=2)

#### Growth study

Figure S7 shows a representative example of how we calculated the growth rate $\mu_{max}$. As example, we describe the growth study of MG1363-GFP $Lac^+$ on 0.09 wt% glucose. We measured the $OD_{660}$ from duplicate shake-flasks hourly (Fig. S7a). We extracted the exponential phase by plotting the log-linear of $OD_{660}$ versus time and fitting a linear curve to the segment where the $\log(OD_{660})$ increased linearly, as determined by the least-squares method (Fig. S7b). We determined the goodness-of-fit for all replicate curves by $R^2$ (Table S2).

#### Gating strategy

Figure S8 shows the flow cytometry gating strategy for a representative example.

**Step 1: Exclude noise** We determined the location of the background noise by the buffer by plotting the forward and side scatter of a sample consisting of only the buffer. This yielded gate 1. From a sample with cells, gate 2 containing everything except the noise was drawn.

**Step 2: Determining the fraction of the fluorescent cheater cells** Gate 2, containing the cells, was the reference for plotting fluorescence against forward scatter. In this plot, the fluorescent cheater cells were marked as Gate 3. To calculate the cheater fraction, we divided the number of these fluorescent cells by the total cell count, excluding the noise.

#### 2 Supplemental figures

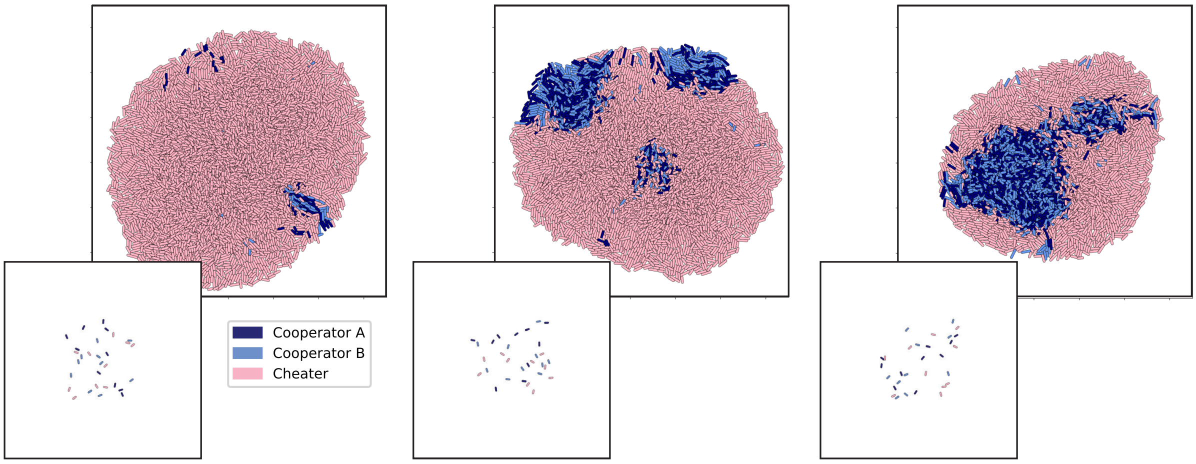


Figure 1: Examples of three different final colonies for the same input parameters for the cross-feeding interaction; value for cost = 0.8 and benefit = 4.0. In the bottom-left corner of each final colony, the initial configuration of that run is shown. Area shown is always 130 × 130 µm$^2$.

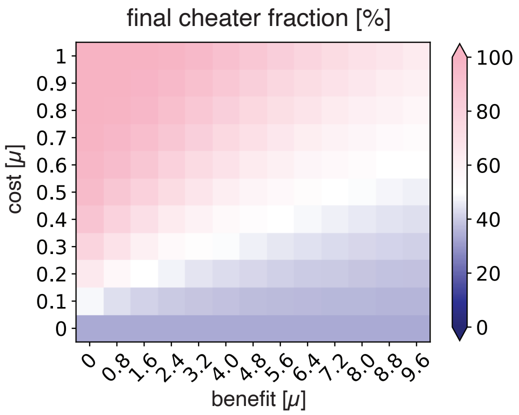


Figure 2: Final cheater fraction for a well-mixed system of A, B and C growing to a population size of $10^4$. Interactions are implemented similarly to the IBM, see SI.

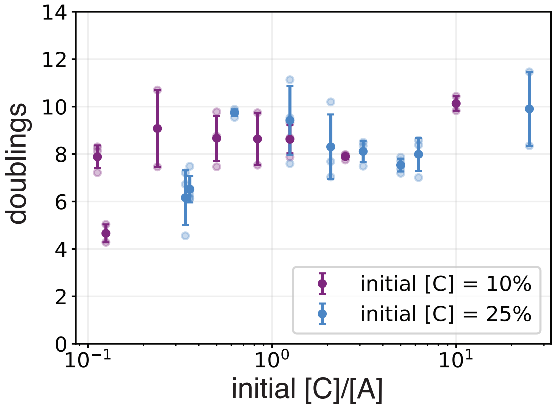


Figure 3: Amount of doublings that occurred on the plates based on the final cell count. The data points correspond to the ones in the cheater fraction curves of Fig. 5a. The corresponding simulation results are based on colonies growing from 30 to 10000 particles which amounts to 8.4 doublings.

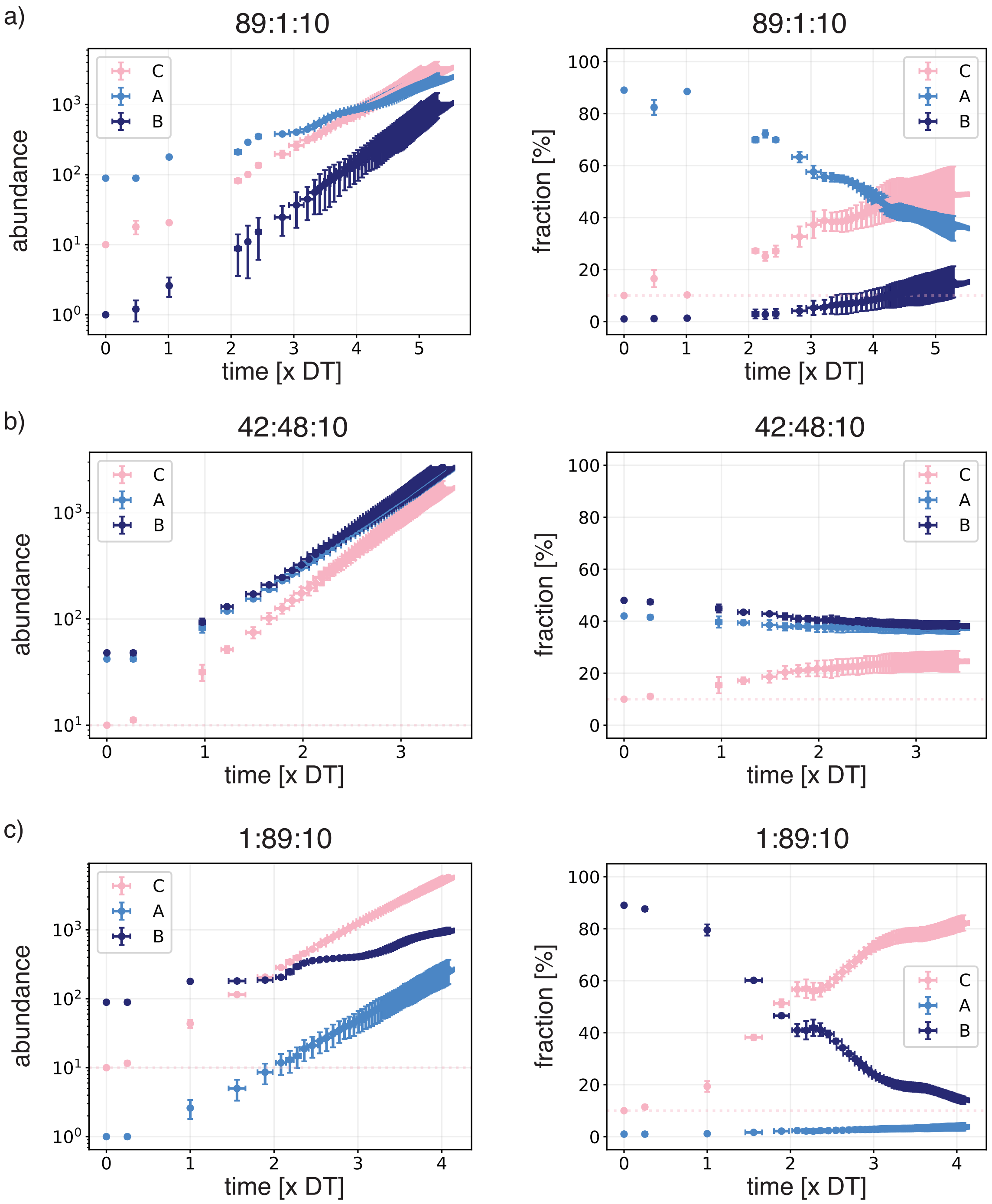


Figure 4: Abundance and fraction of the different particle types over simulation time for the simulation data points of Fig.5b A:B:C ratios a) 89:1:10, b) 42:48:10, and c) 1:89:10. All points are averages and standard deviations of 5 runs.

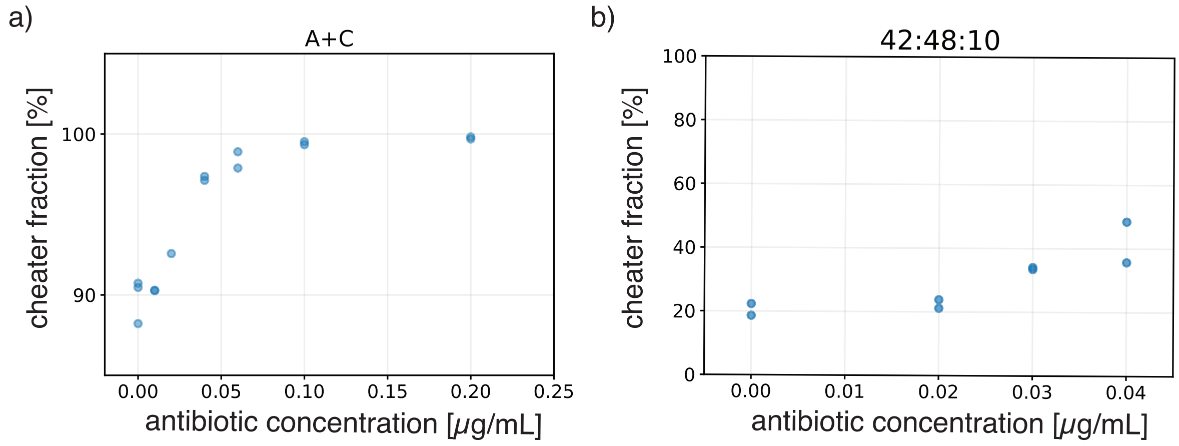


Figure 5: Final cheater fraction for different concentrations of erythromycin a) in plates containing casein and amino acids inoculated with A & C together, and b) in plates containing casein inoculated with A, B & C in a 42:48:10 ratio.

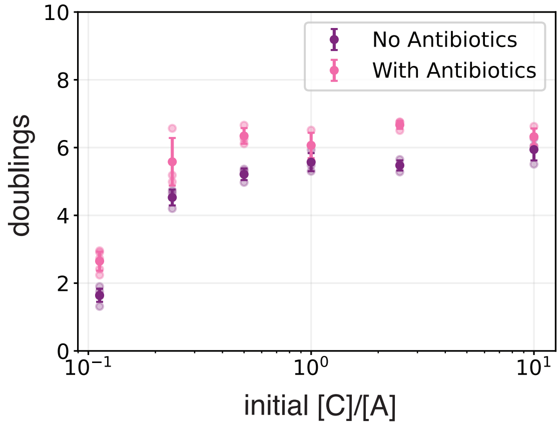


Figure 6: Amount of doublings that occurred on normal plates and plates containing $0.04\,\mu\mathrm{g\,mL^{-1}}$ erythromycin, based on the final cell count. The data points correspond to the ones in the curves of Fig. 6a. The corresponding simulation results are based on colonies growing from 100 to 7000 particles which amounts to 6.1 doublings.

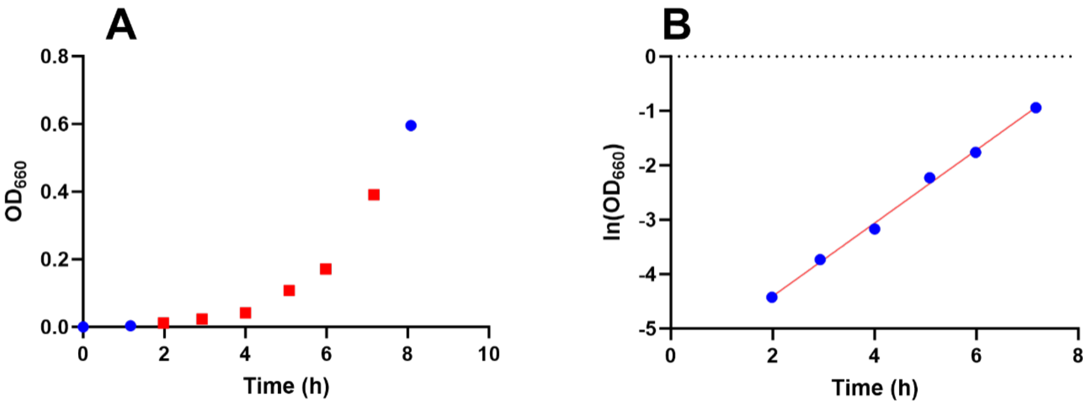


Figure 7: Growth curve of MG1363-GFP Lac$^+$ (cheater). a) $OD_{660}$ over time. The exponential phase is marked with red squares. b) $\ln(OD_{660})$ of the exponential phase measurements over time. A linear curve was fit to the measurements by least-squares fit.

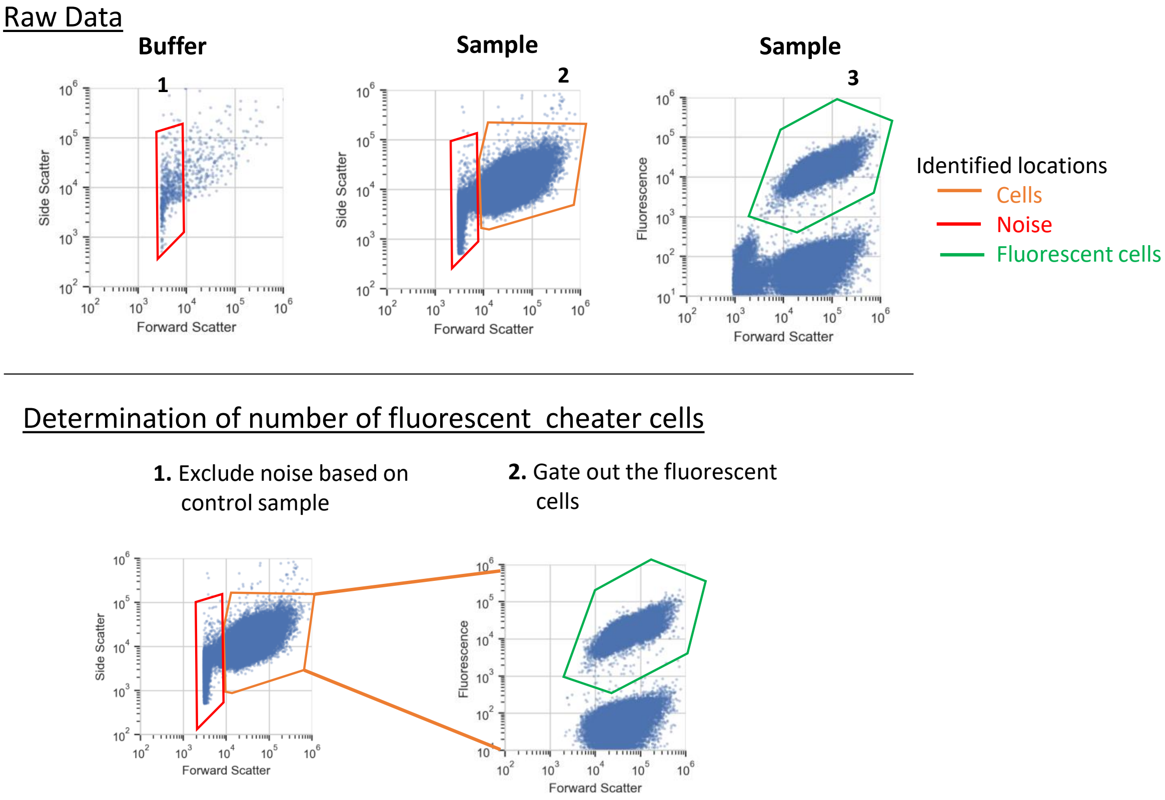


Figure 8: Flow cytometry gating strategy. Areas within the forward scatter – side scatter and fluorescence – forward scatter were identified using control samples. The gating excluded noise and was able to clearly distinguish between fluorescent and non-fluorescent cells to calculate the %cheater fraction.
